# PGAR-Zernike: an ultra-fast, accurate and fully open-source structure retrieval toolkit for convenient structural database construction

**DOI:** 10.1101/2023.03.18.533250

**Authors:** Junhai Qi, Chenjie Feng, Yulin Shi, Jianyi Yang, Fa zhang, Guojun Li, Renmin Han

## Abstract

With the release of AlphaFold2, protein model databases are growing at an unprecedented rate. Efficient structure retrieval schemes are becoming more and more important to quickly analyze structure models. The core problem in structural retrieval is how to measure the similarity between structures. Some structure alignment algorithms can solve this problem but at a substantial time cost. At present, the state-of-the-art method is to convert protein structures into 3D Zernike descriptors and evaluate the similarity between structures by Euclidean distance. However, methods for computing 3D Zernike descriptors of protein structures are almost always based on structural surfaces and most are web servers, which is not conducive for users to analyze customized datasets. To overcome this limitation, we propose PGAR-Zernike, a convenient toolkit for computing different types of Zernike descriptors of structures: the user simply needs to enter one line of command to calculate the Zernike descriptors of all structures in a customized datasets. Compared with the state-of-the-art method based on 3D Zernike descriptors and an efficient structural comparison tool, PGAR-Zernike achieves higher retrieval accuracy and binary classification accuracy on benchmark datasets with different attributes. In addition, we show how PGA-Zernike completes the construction of the descriptor database and the protocol used for the PDB dataset so as to facilitate the local deployment of this tool for interested readers. We construct a demonstration containing 590685 structures; at this scale, our retrieval system takes only 4 ~ 9 seconds to complete a retrieval, and experiments show that it reaches the state-of-the-art level in terms of accuracy. PGAR-Zernike is an open-source toolkit, whose source code and related data are accessible at https://github.com/junhaiqi/PGAR-Zernike/.

## 1 Introduction

Proteins are the basis of all living systems; they fold into specific 3D configurations and perform corresponding biological functions. To understand the mechanism of protein action at the molecular level, it is necessary to accurately predict the 3D structure of protein. In terms of the problem of protein structure prediction, AlphaFold2[1] has achieved comparable prediction accuracy with experimental methods through a well-designed deep neural network and greatly shortened the prediction time, which implies that the protein model structure database will grow at an amazing speed. The latest AlphaFold database release contains over 200 million entries; thus, an efficient and accurate method for measuring protein structural similarity is urgently needed.

Structural alignment is the most direct method to measure the similarity between structures. There are two main types of structural alignment methods: coordinate-based methods and surface-based methods. As early as the 1970s, a scheme[2] was proposed to realize the superimposition of structures based on atomic coordinate information. In the following decades, many similar schemes have been proposed and improved, such as CE[3], DALI[4], RNAAlign[5], TM-align[6], and US-align [7]. Since the nature of the protein surface is crucial to the study of protein-protein (RNA, Ligand) interactions, some structure alignment schemes based on the protein surface have been proposed, such as gmfit[8], ZEAL[9], and ICP[10]. However, these alignment methods are time-consuming. For example, a standard alignment software (gmfit) takes ~0.71 s to complete a structural alignment; it takes ~5 days to complete a structure retrieval (~590,000 alignments).

To measure the similarity between protein structures more efficiently, a common scheme is to convert protein structures into feature vectors, design a metric applied to feature vectors, and convert the problem of measuring the similarity between structures into the problem of measurement calculation between feature vectors. In recent years, two main approaches have been followed to represent structures as feature vectors. One way is to calculate geometric information based on atomic coordinates and convert geometric information into feature vectors. For example, Omokage[11] calculates the distance distribution between feature atoms and proposes the Omokage score to measure the similarity between protein structures. Biozernike’s[12] GEO module converts the geometric information of atoms into a feature vector of 17 lengths. Another way is to convert the protein structure into a protein surface and then calculate the feature vectors based on the protein surface. For example, FTIP[13] extracts 10~90 feature points on the protein surface through the farthest point sampling algorithm and then rapidly calculates the similarity between protein structures based on the feature points and TIPSA algorithm[14]. 3D-SURFER[15, 16, 17] provides a web server for computing 3D Zernike descriptors[18] based on protein surface information (denoted as surface-based 3DZD). Immediately after, [12] proposes an improved scheme for calculating 3D Zernike descriptors based on gmconvert[19].

Zernike descriptors have been widely used not only for retrieval problems but also in other problems of structural biology. For example, interface (binding site) prediction[20, 21, 22, 23, 24], embedding of polymers into EM maps[25], docking problems[26], and analysis of protein surfaces[27]. On the other hand, Zernike descriptors also show their advantages in image reconstruction[28] and 3D structure classification[29]. However, to the best of our knowledge, there is currently no open-source, easy-to-use Zernike implementation for protein (or RNA) structure.

In this study, we propose a toolkit (PGAR-Zernike). One of the unique features of PGAR-Zernike is that it can compute different Zernike descriptors based on different structural representations, such as surface, mesh, and atomic point cloud. Based on this feature, PGAR-Zernike provides two main functions for users: 1) given a customized structural dataset, the descriptor dataset for structural analysis can be constructed; 2) given a query structure, ultra-fast (4~9s) retrieval can be achieved in a large structural dataset [including 590685 structures]. Comprehensive experiments demonstrate that PGAR-Zernike can handle structure retrieval problems efficiently, outperforming surface-based 3DZD and a surface-based alignment tool on benchmark datasets, with a top10-accuracy of ~90% and an AUC value of ~96%. Furthermore, experiments show that the PGAR-based retrieval system (denoted as PGAR-System) is completely feasible for large-scale structural retrieval problems. Given query structures, the top 10~150 structures are output and the average TMscore between them and the query structure is calculated; the average TMscore of the PGAR-System is 0.15~0.2 higher than that of the state-of-the-art retrieval system based on surface-based 3DZD.

## 2 Materials and methods

### 2.1 Overview of PGAR-Zernike

The process of computing the 3D Zernike descriptors of PGAR-Zernike can be divided into the following five steps:

- *Obtain the structural representation*. PGAR-Zernike computes the representation of structures, such as surface, mesh, and atomic point cloud.
- *Extract feature points*. PGAR-Zernike extracts sufficient feature points on the representations of structures that are scaled into the unit sphere.
- *Voxelization*. Based on the feature points, PGAR-Zernike builds a function *f*(*x, y, z*) defined on [-1, 1]^3^ to characterize the structure.
- *Calculate the geometric moments*. Based on the *f*(*x, y, z*), PGAR-Zernike calculates the corresponding geometric moments.
- *Compute descriptors*. The 3D Zernike descriptors are calculated based on specific mathematical expressions.

**Figure 1.** briefly depicts the above five steps. In addition, we improve Omokage[11] and integrate it into the PGAR-Zernike to develop an offline algorithm (called ReOmokage) for computing the geometric features of the structures.

**Figure 1:**
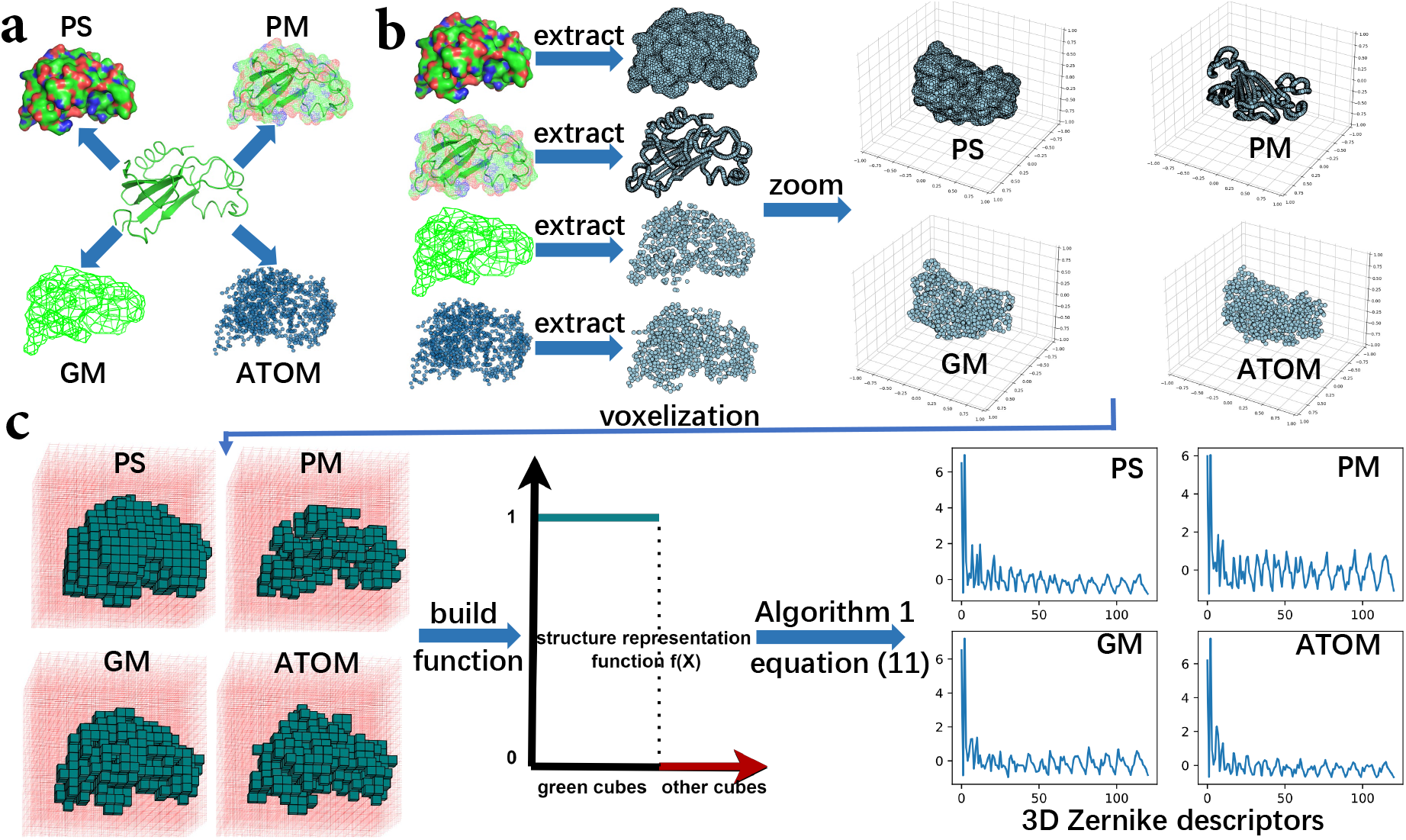
The workflow for computing 3D Zernike descriptors based on the PGAR-Zernike algorithm. **(a)** Given the protein structure(<u>1brnL.pdb</u>), four structural representations in four modes are calculated. **(b)** Feature points are extracted from each structure representation and scaled into unit spheres. **(c)** Based on the feature points, the grid defined at [-1, 1]^3^ is obtained. The green cube is generated by the feature points. The structure representation function *f*(*X*) is defined on the grid and takes values of 1 on the green cubes and 0 on the other cubes. Using Algorithm 1 and Equation 2, four different 3D Zernike descriptors can be computed.

For the problem of measuring structural similarity, ReOmokage outperforms Omokage (refer to **Supplementary Material S3**).

### 2.2 Representation of structure

The core step in computing the 3D Zernike descriptors is to construct an appropriate function *f* (*x, y, z*) to express the structure. The first step in constructing the function *f*(*x, y, z*) is to transform the structure into a reasonable representation. Here, PGAR-Zernike can transform protein structures into multiple forms via <u>pymol</u>[30] and <u>gmconvert</u>[19]. Specifically, PGAR-Zernike can calculate the surface of a protein structure (denoted as PS), mesh (denoted as PM) and GMM-Mesh (denoted as GMM) to represent the structure. In addition, PGAR-Zernike can extract the atomic coordinates of the structure and use these coordinates to represent the structure (denoted as ATOM). All structural representations are shown in **Figure 1a.**.

### 2.3 Extracting feature points

To construct the structure representation function quickly, we first extract feature points from the structure representation. For different structural representations, there are different schemes to extract feature points, and the final number of feature points also varies.

**Figure 2** shows the feature point information and 3D Zernike descriptors of structure (<u>1brnL.pdb</u>) under different representations. As shown in **Figure 2**, the number of feature points generated based on PS is the largest, so these feature points can fully represent the shape of the structure. We can also see that in ATOM and GM modes, the number of feature points is similar and the GMM setting has little effect on feature point extraction. In addition, from **Figure 2c**, we can see that the descriptors in different modes are slightly different, which is caused by the difference in feature points.

**Figure 2:**
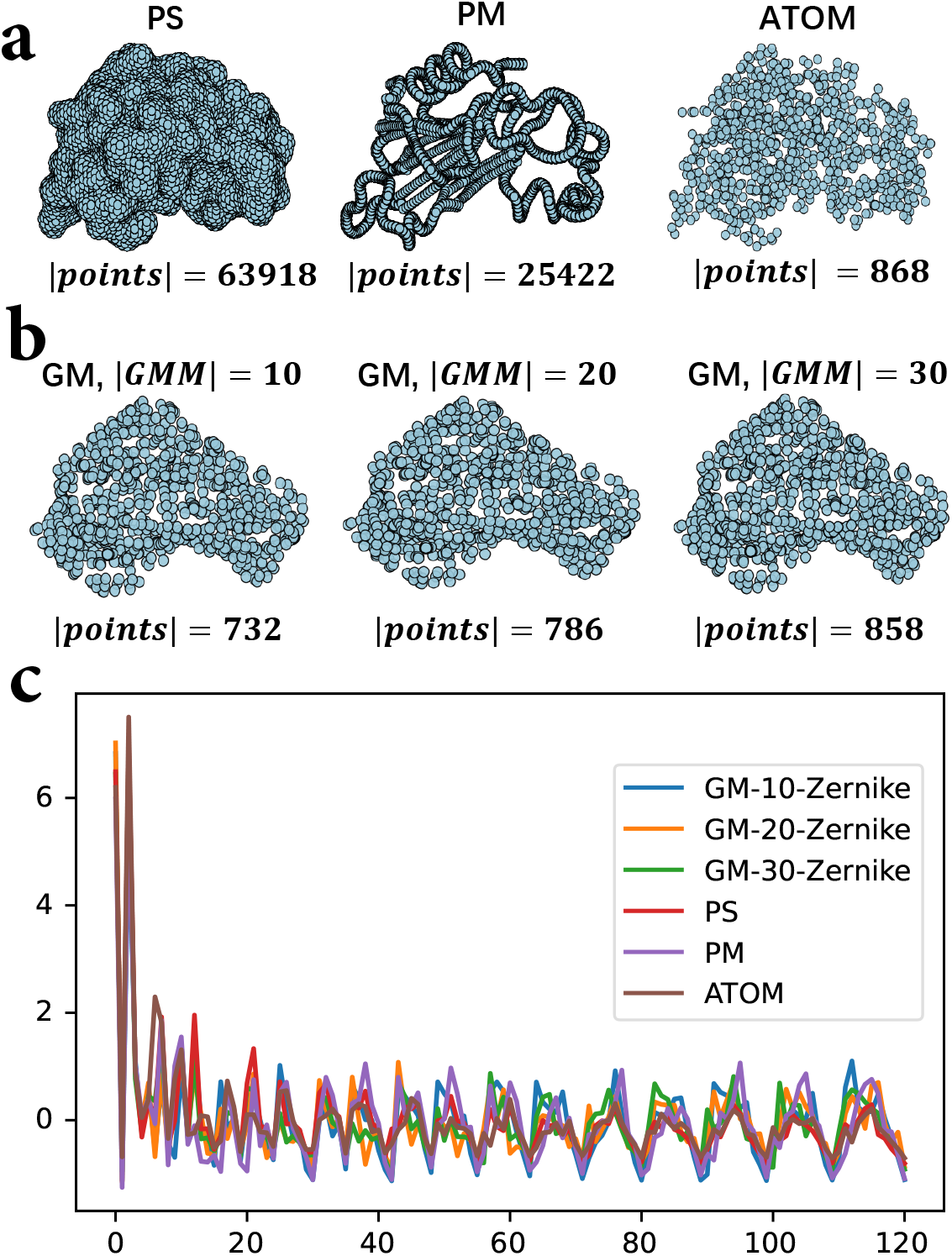
Comparison of the number of feature points and 3D Zernike descriptors in various modes of PGAR-Zernike. **(a)** The shape and quantity of feature points in PS, PM, and ATOM modes. **(b)** For different GMM parameters, the shape and number of feature points in GM mode. **(c)** Six 3D Zernike descriptors in six modes.

The feature points from different schemes have different properties. For example, although a large number of feature points can be extracted based on the surface of the structure, these feature points cannot express the internal information of the structure, while the feature points extracted based on atomic points can express such information. For different structural biology problems, different feature points may be needed to express the structure, so our method provides different schemes to calculate the feature points of the structure.

### 2.4 Structure representation function

All the feature points are embedded in the unit sphere. Let the set of feature points be *S*, *∀*(*x, y, z*) ∈ *S*, it satisfies |*x*|^2^ + |*y*|^2^ + |*z*|^2^ ≤ 1. In addition, we build a grid (*N* × *N* × *N*, default *N* = 200) defined on [-1, 1]^3^ with a total of *N*^3^ cells, and the set of cells is defined as *C* = {*cell*_(0,0,0)_, *cell*_(1,0,0)_,…, *cell*_(N-1,N-1,N-1)_ }. Equation (1) is a mathematical expression for the structural representation function. When *S* is given, the computational complexity of *f*(*x, y, z*) is *O*(|*S*|) (refer to Algorithm 2).

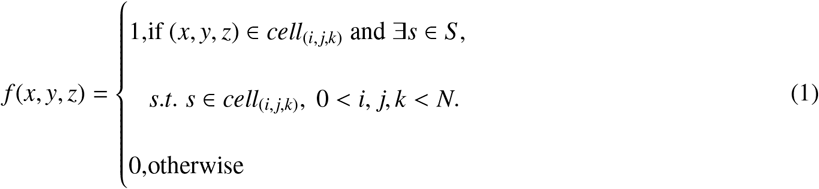

#### Algorithm 1 PGAR-Zernike algorithm for computing geometric moments.

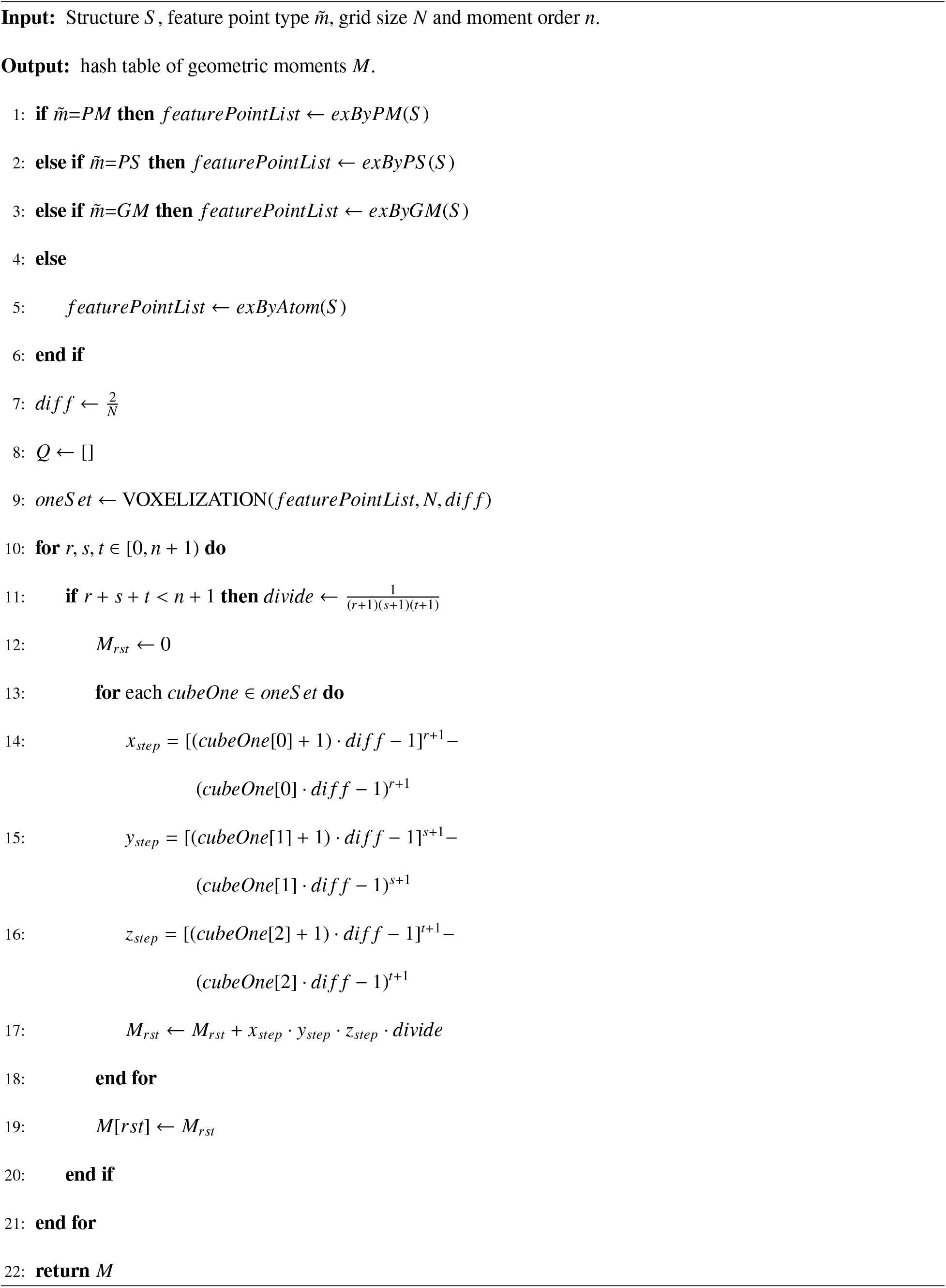

#### Algorithm 2 PGAR-Zernike algorithm for building the structure representation function.

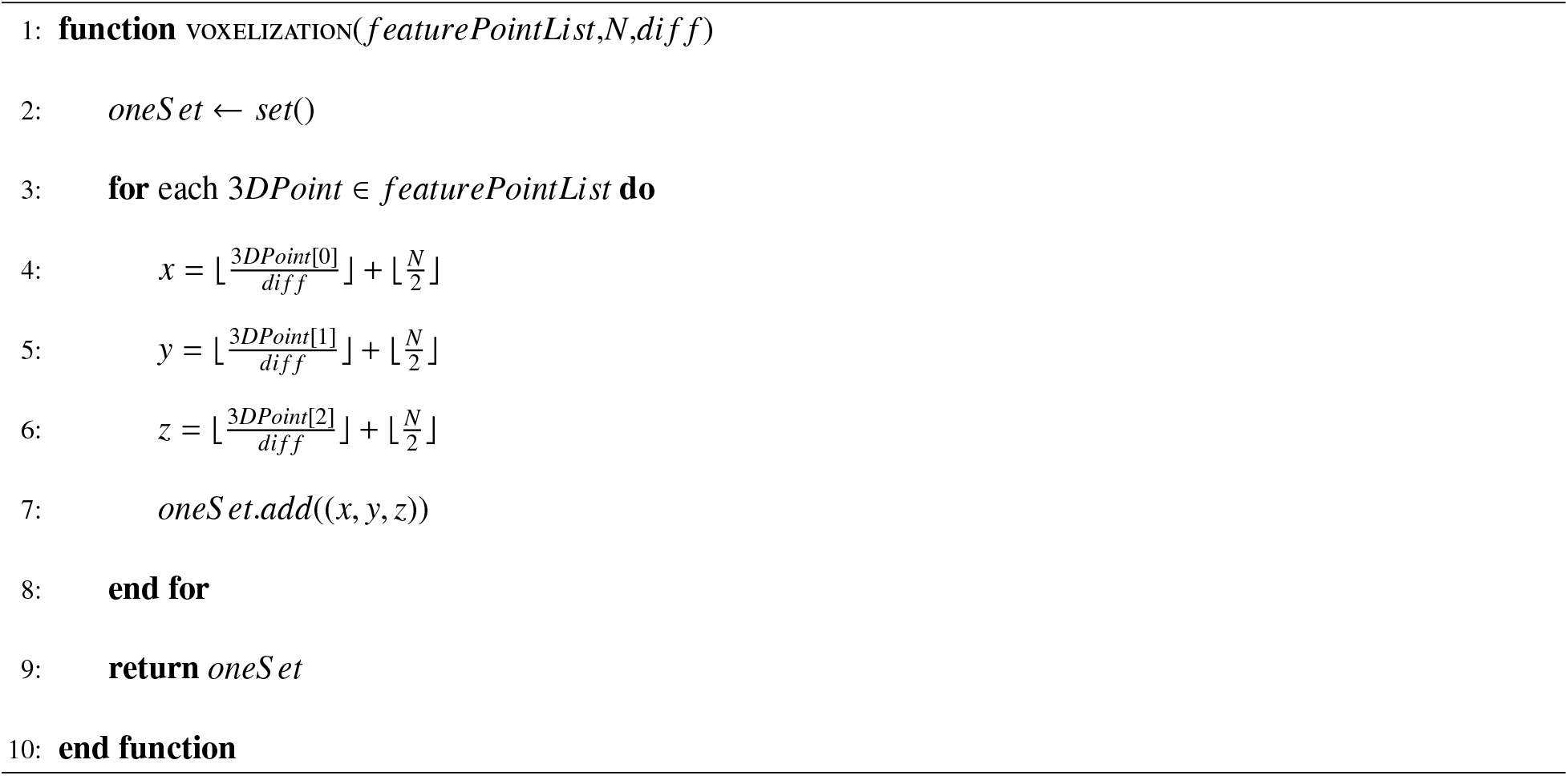

### 2.5 PGAR-Zernike for computing 3D Zernike descriptors

Computing the 3D Zernike descriptor essentially involves computing 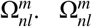 can be calculated using Equation (2)[18], where the detailed analysis is given in **supplementary material S1**.

From Equation (2) ~ (4), we can see that 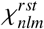 and *f*(*X*) are independent, and 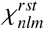 can be computed when order n is given. Therefore, the key to calculating the 3D Zernike descriptor is to construct *f*(*X*) and calculate *M_rst_* (geometric moments). Here, the definitions of 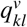 and 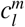 are given in the **supplementary material S1.3**.

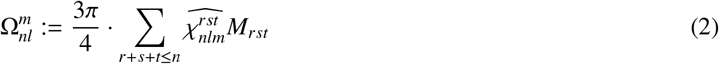

Here, 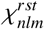 is defined in Equation (3), and *M_rst_* is defined in Equation (4).

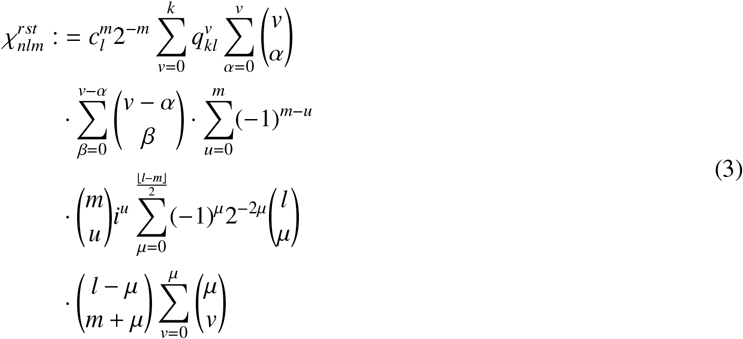

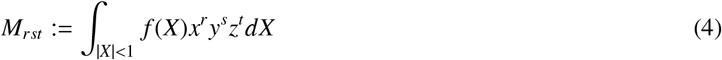

Based on previous work[18] and the definition of an integral, we can derive the discrete form of Equation (4), namely, Equation (5).

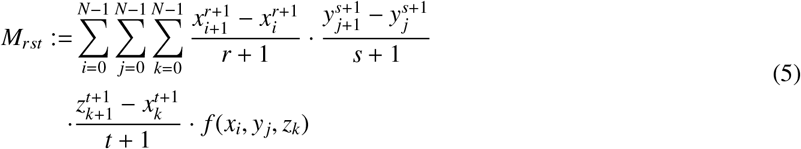

Note that we define the structure representation function *f*(*X*) on the interval [-1,1]^3^. For *∀_s_* ∈ *S*, we have |*s*| < 1.

Let *cube* = {(*x, y, z*)| |*x*|, |*y*|, |*z*| ≤ 1} and *unitBall* = {(*x, y, z*)| *x*^2^ + *y*^2^ + *z*^2^ ≤ 1}; the following formula holds:

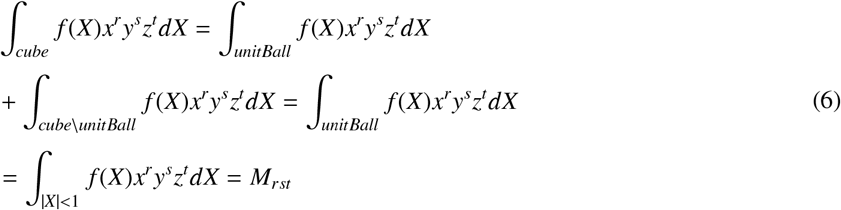

Therefore, the integral of *f*(*X*) on *cube* is equivalent to the integral of *f*(*X*) on *unitBall*.

Algorithm 1 shows the process of computing geometric moments. Initially, Algorithm 1 calculates the feature points of the structure[lines 1-6] based on the modes input by the user. There are a total of four main functions, namely, *exByPM, exByPS*, *exByGM*, and *exByAtom*, and the details of their calculations can be found in **Supplementary Material S2.1**. Then, Algorithm 1 determines the size of each small cube in the grid according to the size of the grid[line 7] and calculates the small cubes whose structure representation function takes a value of 1[line 9]. Finally, the geometric moments are calculated according to Equation (5)[lines 10-21].

Algorithm 2 shows the process of constructing a structural representation function *f*(*X*). Algorithm 2 determines the indexes of the small cubes containing the feature points: *f*(*X*) takes a value of 1 in these small cubes, and takes a value of 0 in other cubes.

After all geometric moments are calculated, the 3D Zernike descriptors can be obtained via Equation (2) ~ Equation (4).

### 2.6 Build the descriptor dataset based on PGAR-Zernike

A convenient script has been integrated into PGAR-Zernike to enable users to quickly build descriptor datasets. For a folder that contains only structures (<u>pdb</u> format), the user needs to execute only the one-line command to calculate the descriptors of all structures, which implies that PGAR-Zernike can be easily integrated into other structural analysis algorithms. The efficiency of building the descriptor dataset is related to the complexity of the structure. For more details, please refer to the **Supplementary Material S4.3**.

### 2.7 PGAR-System

We successfully downloaded the entire PDB database (https://www.rcsb.org/), which contains 193,728 protein structures. We split all the structures into single-stranded structures, obtained a large database of 590,685 structures, and calculated the descriptors (PM mode) of each structure to construct the retrieval database. Based on this retrieval database, we built a retrieval system (denoted as PGAR-System). The following is the workflow of the PGAR-System. Given the query structure, we first determine whether it is in the database. If not, we calculate its descriptor (denoted by query descriptor), then calculate the Euclidean distance between all descriptors in the database and the query descriptor, and finally rank the results according to the Euclidean distance (ascending order). With the default parameters, the user obtains the top 100 structures.

### 2.8 Usage of PGAR-Zernike

For some main functions delivered by PGAR-Zernike for users, namely, the calculation of different types of descriptors, the construction of the descriptor database and the use of PGAR-System, there are detailed usage instructions in the **Supplementary Material S5**.

## 3 Results

### 3.1 Benchmarking

#### 3.1.1 Benchmark data sets

We carefully constructed benchmark datasets based on the basic local alignment search tool[31], TM-align[6] and web server[32]. To initially test the performance of PGAR-Zernike, we constructed two protein structure datasets and one RNA structure dataset, denoted as **Protein160**, **Protein13** and **RNA16**. Each dataset has different characteristics, and the details of these datasets are presented in **Table 2**. In addition, to test the binary classification ability of PGAR-Zernike, we obtained the set of structure pairs of the above three datasets. All the structure pairs were divided into positive samples and negative samples, and the detailed information is provided in **Table 3**. To test the performance of PGAR-System, we randomly selected 500 structures from the entire single-stranded structure database to form a dataset (denoted as **Random500**). All details of constructing the datasets are provided in **Supplementary Material S2.2**.

**Table 1:** The top-*k* accuracy of our method and reference methods on the benchmark data sets. For each row in the table, the best is underlined, and the next best is double underlined.

|  |  | PGAR-Zernike components |  |  |  |  | Reference methods |  |
| --- | --- | --- | --- | --- | --- | --- | --- | --- |
| Data set | top-$k$ | PM-Zernike | PS-Zernike | GM-Zernike | ATOM-Zernike | ReOmokage | 3D-SURFER | gmfit |
| Protein160 | top-10 | <u>0.9114</u> | 0.854 | 0.6015 | 0.8378 | 0.5624 | 0.8762 | <u>0.9093</u> |
|  | top-30 | <u>0.9463</u> | 0.8952 | 0.7148 | 0.8842 | 0.7364 | 0.9103 | <u>0.9237</u> |
|  | top-50 | <u>0.954</u> | 0.916 | 0.7663 | 0.9026 | 0.8022 | 0.9238 | <u>0.9309</u> |
|  | top-70 | <u>0.9571</u> | 0.9257 | 0.7916 | 0.9109 | 0.8438 | 0.932 | <u>0.9341</u> |
|  | top-100 | <u>0.9596</u> | 0.9365 | 0.82 | 0.9299 | 0.8761 | <u>0.9465</u> | 0.946 |
| Average accuracy |  | <u>0.94568</u> | 0.90548 | 0.73884 | 0.89308 | 0.76418 | 0.91816 | <u>0.9288</u> |
| Protein13 | top-10 | <u>1</u> | 1 | 0.9615 | 1 | 1 | — | <u>1</u> |
|  | top-30 | 0.9673 | 0.9628 | 0.9393 | <u>0.9825</u> | 0.9673 | — | <u>0.9912</u> |
|  | top-50 | 0.9642 | 0.9592 | 0.9452 | <u>0.988</u> | 0.9627 | — | <u>0.9672</u> |
|  | top-70 | 0.9431 | 0.9284 | 0.9207 | <u>0.9864</u> | 0.942 | — | <u>0.9536</u> |
|  | top-100 | 0.9288 | 0.9304 | 0.9183 | <u>0.9899</u> | 0.9264 | — | <u>0.9451</u> |
| Average accuracy |  | 0.96068 | 0.95616 | 0.937 | <u>0.98936</u> | 0.95968 | — | <u>0.97142</u> |
| RNA16 | top-10 | 0.8919 | <u>0.92</u> | <u>0.9263</u> | 0.8441 | 0.9109 | — | 0.8326 |
|  | top-30 | 0.8982 | <u>0.9201</u> | 0.9017 | 0.8818 | <u>0.9181</u> | — | 0.7571 |
|  | top-50 | <u>0.91</u> | <u>0.9198</u> | 0.899 | 0.9097 | 0.9098 | — | 0.7495 |
|  | top-70 | <u>0.9067</u> | <u>0.9394</u> | 0.9054 | 0.9066 | 0.9053 | — | 0.744 |
|  | top-100 | <u>0.9359</u> | <u>0.9486</u> | 0.9246 | 0.9075 | 0.9081 | — | 0.7442 |
| Average accuracy |  | 0.90854 | <u>0.92958</u> | <u>0.9114</u> | 0.88994 | 0.91044 | — | 0.76548 |

**Table 2:**
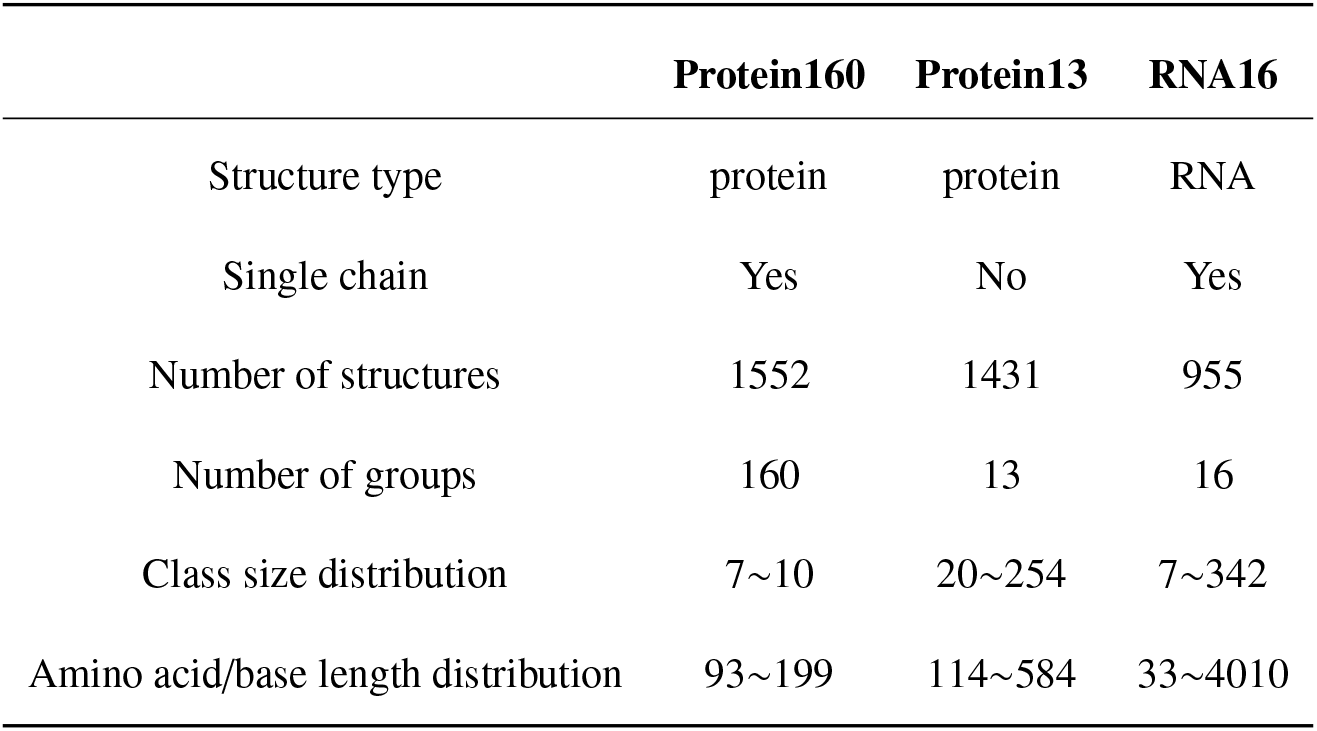
Details about **Protein160**, **Protein13** and **RNA16**.

**Table 3:** Details about **Protein160-Pairs**, **Protein13-Pairs** and **RNA16-Pairs**.

|  | **Protein160-Pairs** | **Protein13-Pairs** | **RNA16-Pairs** |
| --- | --- | --- | --- |
| Number of positive samples | 1552 | 1431 | 955 |
| Number of negative samples | 246768 | 17172 | 14325 |

#### 3.1.2 Reference methods

Our method is compared with 3D-SURFER and gmfit[8]. Currently, 3D-SURFER is the state-of-the-art protein structure retrieval system [include 606,272 structure chains] based on surface-based 3DZD, so we compare our method with it. gmfit can quickly complete the structure comparison on the Gaussian mixture model of the structures, which means that users can calculate the Gaussian mixture model database in advance and then build a retrieval system based on gmfit; thus, our method is also compared with gmfit. Please refer to **Supplementary Material S2.3** for details on the use of 3D-SURFER and gmfit.

#### 3.1.3 Experimental environment

All the experiments were run on an Ubuntu 20.04 system with Intel(R) Core(TM) i9-10980XE (18 cores), 128 GB memory, and an NVIDIA RTX3080.

### 3.2 Search procedure and evaluation

#### 3.2.1 Initial evaluation on benchmark datasets

For a given structure, PGAR-Zernike can compute four different Zernike descriptors based on four different representations. To decide which descriptor is best for structure retrieval, we design experiments based on benchmark datasets. The details about the evaluation metric are given in the **Supplementary Material S4.1**.

We successfully obtain the descriptors of **Protein160** through 3D-SURFER’s web server. However, we obtain 3D Zernike descriptors for fewer than 1000 structures in **Protein13** using 3D-SURFER, and 3D-SURFER is not suitable for computing descriptors of RNA structures. Therefore, our method is not compared with 3D-SURFER on **Protein13** and **RNA16**.

**Table 1** shows the top-k accuracy of our method and the two reference methods on **Protein160**, **Protein13**, and **RNA16**. As shown in **Table 1**, among the five modules of PGAR-Zernike, PM-Zernike is most stable and achieves better results on three datasets, slightly ahead of gmfit and 3D-SURFER. The top-10 accuracy of GM-Zernike and ReOmokage is slightly worse on **Protein160**, but it is outstanding on **RNA16**, probably because GM-Zernike and ReOmokage do not extract sufficient feature points and ignore the internal structure information; thus, they perform slightly worse on datasets containing complex structures. All methods have high top-k accuracy on **Protein13** due to the small number of groups in **Protein13** and the criterion that two structures belong to the same group is more stringent, which makes the retrieval problem less difficult. In particular, the top-k accuracy of PGAR-Zernike on **RNA16** is relatively good compared with that of gmfit, indicating that PGAR-Zernike is more suitable for measuring the similarity between RNA structures.

On the other hand, we evaluate the binary classification performance of PGAR-Zernike on **Structure-Pairs**. **Figure 3** shows the receiver operating characteristic (ROC) and precision-recall (PR) curves of all methods. Overall, the binary classification accuracy of all methods is basically consistent with their top-*k* accuracy.

**Figure 3:**
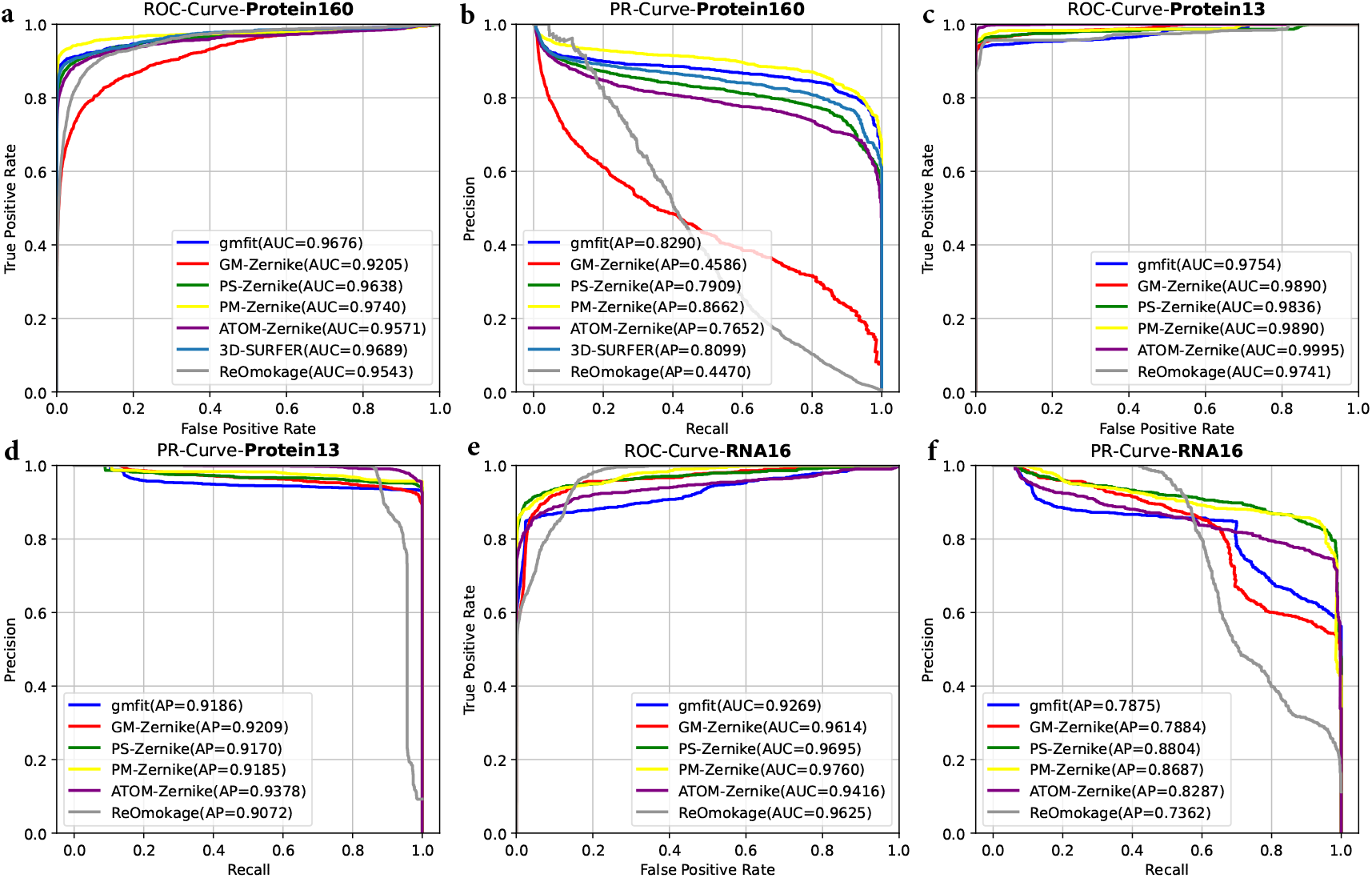
Structure retrieval performance comparison of PGAR-Zernike, 3D-SURFER and gmfit. (**a**) ROC curves and AUC values, (**b**) PR curves and AP values of our method and the reference methods on **Protein160-Pairs**. (**c**) ROC curves and AUC values, (**d**) PR curves and AP values of our method and the reference methods on **Protein13-Pairs**. (**e**) ROC curves and AUC values, (**f**) PR curves and AP values of our method and the reference methods on **RNA16-Pairs**.

The above experiments show that PM-Zernike achieves the best performance, so we use it to build a retrieval system (PGAR-System).

#### 3.2.2 Performance of PGAR-System

We take each structure in **Random500** as a query structure and input 3D-SURFER and PGAR-System to obtain the retrieval results. Here, we use usalign[7] to calculate the TMscore and the RMSD between the structure in the retrieval results and the query structure and use these two metrics to evaluate the performance of the retrieval system. **Table 4** presents the main results. From **Table 4**, we can see that the average TMscore of PGAR-Zernike is 0.15~0.2 higher than that of 3D-SURFER, and the average RMSD is ~1Å lower than that of 3D-SURFER. The results indicate that PM-Zernike is more suitable than surface-based 3DZD for characterizing the structure.

**Table 4:** Retrieval evaluation of 3D-SURFER and PGAR-System on **Random500**. All values in the table (mean±standard deviation) are calculated by usalign[7].

|  |  | top-10 | top-30 | top-50 | top-100 | top-150 |
| --- | --- | --- | --- | --- | --- | --- |
| 3D-SURFER | TMscore | 0.606±0.357 | 0.548±0.350 | 0.522±0.343 | 0.486±0.331 | 0.464±0.320 |
| | RMSD($\text{\AA}$) | 2.698±2.253 | 3.133±2.269 | 3.328±2.252 | 3.595±2.217 | 3.742±2.172 |
| PGAR-System | TMscore | 0.835±0.269 | 0.751±0.314 | 0.708±0.328 | 0.650±0.337 | 0.614±0.339 |
| | RMSD($\text{\AA}$) | 1.46±1.879 | 2.086±2.221 | 2.386±2.326 | 2.812±2.428 | 3.067±2.465 |

#### 3.2.3 Speed of PGAR-System

In the entire structure database, we randomly selected 1000 structure pairs, used TMalign, DeepAlign[33], usalign, gmfit and PGAR-Zernike to complete the alignment of 1000 structure pairs, and obtained the average time; Table 5 shows the main results. As shown in Table 5, PGAR-Zernike has a very obvious efficiency advantage compared with other structural alignment tools. The main reason is that PGAR-Zernike measures the structural similarity by calculating the Euclidean distance between the descriptors of the structure, and the length of the descriptor is fixed, which means the time complexity of measuring the structural similarity is O(1).

**Table 5:** Average running time for the five tools to compute a structural similarity.

|  | PGAR-Zernike | gmfit | TMalign | DeepAlign | USalign |
| --- | --- | --- | --- | --- | --- |
| run time(s) | $2.75 \times 10^{-6}$ | 0.710 | 0.038 | 0.221 | 0.078 |

Given the query structure, there are two cases: (1) it is in our database or (2) it is not in our database. If (1) is established, it takes ~4 s for PGAR-System to complete the entire retrieval process. If (2) holds, PGAR-System needs to first calculate the descriptor of the query structure. For a medium-sized structure (~300 amino acids), it takes approximately ~5 s to complete the computation of the descriptor [PM mode]. Therefore, if (2) holds, it takes ~9 s for PGAR-System to complete the entire retrieval process.

### 3.3 Statistical significance

We completed the significance experiment to facilitate a preliminary estimate of the similarity between structures based on Euclidean distance. Since the above experiments show that GM-Zernike cannot well characterize the structure, we do not perform corresponding experiments for GM-Zernike.

The statistical significance of PGAR-Zernike-Euclidean-distance is estimated by comparing approximately 1.99 million random structure pairs from our downloaded protein structure database (193,728 entries). A list of P-values and corresponding PGAR-ATOM/PGAR-PM/PGAR-PS is given in **Table 6**. For example, a PGAR-ATOM/PGAR-PM/PGAR-PS of 2.44/3.93/2.77 indicates significant similarity at a P-value of 0.01. One may use these distance values for different Zernike descriptors to quickly estimate the similarity between structures.

**Table 6:** Statistical significance of the Euclidean distance between PGAR-Zernike descriptors of structures.

| P-value | $5 \times 10^{-2}$ | $1 \times 10^{-2}$ | $1 \times 10^{-3}$ | $1 \times 10^{-4}$ | $1 \times 10^{-5}$ | $1 \times 10^{-6}$ |
| --- | --- | --- | --- | --- | --- | --- |
| PGAR-ATOM | 2.987906 | 2.442935 | 1.988947 | 1.477029 | 0.522746 | 0.080739 |
| PGAR-PM | 4.653991 | 3.932048 | 3.169391 | 1.259989 | 0.252537 | 0.000244 |
| PGAR-PS | 3.299481 | 2.772549 | 2.306926 | 1.411444 | 0.514943 | 0.285697 |

## 4 Discussion

With the rapid growth of protein structure databases, an efficient and accurate structural similarity analysis scheme is urgently needed. Existing structure alignment algorithms cannot efficiently analyze the similarity between structures. In this work, we propose PGAR-Zernike, which allows users to compute different types of 3D Zernike descriptors and geometric features. Based on PGAR-Zernike, we build a retrieval system[PGAR-System] and demonstrate that it out-performs the state-of-the-art retrieval system based on surface-based 3DZD. In terms of efficiency, PGAR-System can complete a retrieval in less than 10 seconds. In particular, users can easily deploy PGAR-Zernike in the local system for structure retrieval and descriptor database construction and integrate into other structure analysis algorithms.

